# Broadband Rheometry of Native Soft Tissues via i-Rheo-Indent

**DOI:** 10.64898/2026.10.23.761029

**Authors:** Andrea Jannina Fernandez, Laura Jenkins, Graham M. Gibson, Manlio Tassieri

## Abstract

Characterising the frequency-dependent viscoelasticity of soft biological tissues is important for mechanobiology, disease phenotyping, and the design of biomimetic materials. However, conventional oscillatory rheometry becomes increasingly compromised by instrumental inertia at high frequencies and often requires specimens to be cut, reshaped, or compressed to conform to predefined measuring geometries, while local indentation methods probe only limited tissue volumes. Here, we apply *i-Rheo-Indent*, a macroscale indentation approach that transforms the measured force-relaxation and indentation histories from a single experiment directly into the broadband viscoelastic spectrum, without fitting to a predefined constitutive model. Measurements on a 3D-printed porcine liver analogue and murine quadriceps and brain showed good agreement with conventional oscillatory rheometry over the frequency range in which the latter remained reliable, while extending the accessible response up to 500 rad, s^*−*1^, beyond the regime increasingly dominated by instrument inertia. The method also enabled direct characterisation of intact tubular murine colon, yielding an elastic response consistent with published indentation measurements while additionally resolving the frequency-dependent viscous contribution. Intact murine livers exhibited substantially larger viscoelastic moduli than biopsy-punched specimens prepared for parallel-plate rheometry, indicating that specimen preparation can materially influence the measured mechanical response. By combining rapid broadband acquisition, macroscale probing, minimal specimen manipulation, and comparatively simple instrumentation, *i-Rheo-Indent* provides a practical route to characterising soft tissues closer to their native mechanical state, with potential applications in mechanobiology, disease phenotyping, and the development of tissue-engineered and biomimetic materials.

## 1 Introduction

Over the past two decades, the mechanical microenvironment of cells has progressively emerged from being regarded as a passive structural support to being recognised as an active regulator of biological function. Landmark studies in the mid-2000s demonstrated that cells sense and respond to the mechanical properties of their surroundings, and that substrate stiffness can directly influence cell morphology, adhesion, migration and lineage commitment [1, 2, 3]. This recognition has had profound implications across mechanobiology, tissue engineering and regenerative medicine. Changes in tissue stiffness have subsequently been associated with ageing [4, 5, 6] and disease progression [7, 8, 9], while mechanical properties are increasingly being investigated as potential biomarkers for diagnostic applications [10, 11, 12]. In parallel, the development of tissue-mimicking hydrogels and three-dimensional scaffolds has increasingly relied on reproducing not only the biochemical but also the mechanical environment of native extracellular matrix (ECM) [13].

Importantly, however, biological tissues and their synthetic analogues are not purely elastic materials—they are viscoelastic and display time-dependent relaxation and energy dissipation. Consequently, a single measure of “stiffness” cannot fully describe the mechanical environment experienced by cells. Indeed, recent studies have demonstrated that the viscous component of otherwise visoelastic matrices can itself regulate mechanotransduction and cell fate, while matrix stress-relaxation dynamics can strongly influence cellular behaviour [14, 15]. A meaningful mechanical characterisation of biological tissues therefore requires access to their frequency-dependent linear viscoelastic response, described by the complex shear modulus *G*^*\**^(*ω*) = *G*^*′*^(*ω*)+ *iG*^*′′*^(*ω*), whose storage and loss components quantify the elastic and dissipative contributions, respectively.

Oscillatory rotational rheometry remains the gold standard for determining these frequency-dependent properties at the bulk scale [16]. Nevertheless, its application to soft biological tissues presents a number of fundamental experimental challenges. Unlike liquids, soft solids do not naturally conform to the geometry of the measuring tools, and sufficient sample–tool contact must therefore be established before a reliable torque can be measured. This commonly requires compression of the specimen during loading. Such compression is not necessarily mechanically innocuous: biomimetic hydrogels have been shown to exhibit pronounced compressional stress stiffening or softening, meaning that the normal force applied simply to establish contact can modify the mechanical state that the experiment is intended to measure [17, 18]. The problem becomes particularly relevant when small-diameter tools are used to accommodate the limited volume of biological specimens, since a given normal force then produces a larger compressional stress. Interfacial phenomena such as wall slip and sample debonding provide additional sources of uncertainty [19]. Moreover, tissue biopsies and excised organs are often small, heterogeneous and irregularly shaped, and may need to be cut, punched or otherwise reshaped before they can be accommodated between conventional parallel plates. The preparation required to make a biological specimen compatible with the measurement geometry may therefore alter the very mechanical properties being investigated.

A further limitation arises at high frequencies from the inertia of the measuring system. As discussed comprehensively by Ewoldt *et al*. [20], reliable oscillatory measurements require the torque generated by the material to remain sufficiently large relative to the instrumental inertial contribution. This condition becomes especially difficult to satisfy for compliant materials such as soft tissues. Once instrument inertia becomes significant, the measured moduli can develop an apparent high-frequency response approaching an *ω*^2^ dependence and may be mistakenly interpreted as an intrinsic material property (as illustrated in Figures 1C and 1F). Related inertio-elastic ringing can also contaminate the short-time response of step measurements [20]. Although instrumental corrections can extend the accessible range, the resulting data require particular care once the sample response approaches the instrumental limit.

**Figure 1:**
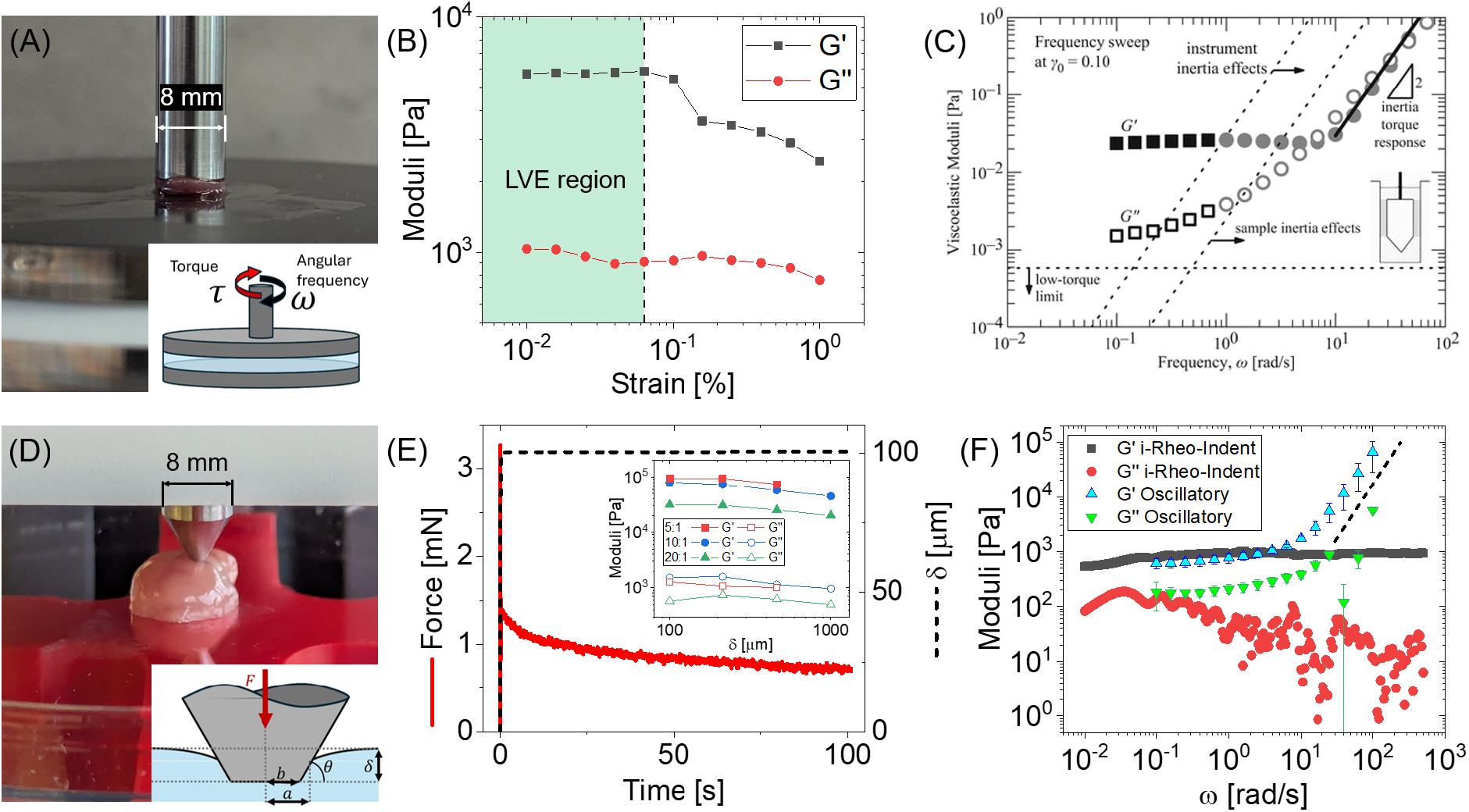
Principle and experimental implementation of conventional linear oscillatory rheometry and the *i-Rheo-Indent* methodology [35] for soft biological tissues. **(A)** Conventional parallel-plate oscillatory rheometry on cut liver using an 8-mm measuring geometry. The schematic illustrates the oscillatory torque *T* applied by the rheometer at angular frequency *ω*, from which the corresponding shear stress *τ* is determined. Reproduced from [35]. **(B)** Representative strain sweep obtained from murine quadriceps tissue at *ω* = 10 rad, s^*−*1^. The shaded green region identifies the linear viscoelastic (LVE) regime, over which the measured moduli remain approximately independent of strain; the vertical dashed line marks its approximate upper limit. **(C)** Low-torque and inertial limits for an oscillatory frequency sweep of hagfish gel, illustrating the frequency range over which instrumental effects can dominate the measured response. The low-torque, instrument-inertia and sample-inertia limits are shown together with the predicted inertial torque response for an instrument-inertia correction error of *ε* = 0.01. *Gray circles* indicate the abrupt change in the raw phase angle from *<* 15^*°*^ to *>* 130^*°*^, providing an additional indication that instrument-inertia corrections become necessary. Data originally published in [37]; reproduced from [20] with permission from Springer Nature. **(D)** Experimental set-up used for *i-Rheo-Indent*, consisting of a Mecmesin OmniTest 2.5 tensile/compression testing machine equipped with a 2-N load cell and a truncated-conical probe (base diameter: 8 mm; blunt-end diameter: 0.5 mm; cone angle: 60^*°*^). A wild-type murine brain is shown being indented at the cerebrum to a depth of 100 *µ*m. The inset illustrates the truncated-conical indentation geometry and its defining parameters, including indentation depth *δ*, contact radius *a*, blunt-end radius *b*, and cone angle *θ*. Reproduced from [35]. **(E)** Representative force and displacement histories obtained from the murine brain measurement. The imposed displacement (black dashed line) is rapidly increased and maintained at approximately 100 *µ*m, while the corresponding normal force (red) undergoes viscoelastic relaxation. These signals are transformed according to Eqs. 3 and 5 to recover the frequency-dependent viscoelastic response. The inset illustrates the dependence of the recovered moduli on indentation depth for cured polydimethylsiloxane (PDMS) samples with base-to-curing-agent ratios of 5:1, 10:1, and 20:1, providing an indentation analogue of an amplitude sweep for identifying the range over which the recovered moduli remain independent of the imposed deformation; values represent averages over 0.1–1 rad, s^*−*1^ from individual indentation measurements. **(F)** Viscoelastic spectra recovered from the force– indentation data in (E), compared with conventional oscillatory measurements on murine brain tissue (*n* = 3). The black dashed line indicates the approximate *ω*^2^ scaling characteristic of the high-frequency instrumental-inertia artefact in the oscillatory measurements.

These difficulties have motivated the widespread adoption of indentation-based and microme-chanical approaches, particularly atomic force microscopy (AFM), for the characterisation of biological materials. AFM has been applied from tissues and organs [21, 22, 23] down to individual cells [24, 25, 26, 27], offering exceptionally high spatial resolution while requiring only small specimens. This spatial resolution, however, also represents a limitation when the objective is to determine representative bulk tissue properties. The very small region interrogated by an AFM probe makes the measurement intrinsically sensitive to local structural heterogeneity [28, 29, 30], and the measured modulus can depend strongly on probe size and geometry [31]. Moreover, extraction of mechanical properties generally relies on fitting force–indentation data using contact-mechanics descriptions, commonly Hertz- or Sneddon-type formulations and their extensions. These require assumptions concerning probe geometry, sample homogeneity, Poisson’s ratio, contact conditions and deformation regime; departures from such assumptions can introduce systematic differences in the recovered modulus [32, 33]. Consequently, AFM-derived nanomechanical properties need not coincide with the macroscale response of a heterogeneous tissue [34]. AFM measurements also require specialised instrumentation and considerable experimental expertise, limiting their accessibility for routine rheological characterisation.

Here, we bridge the gap between conventional bulk rheometry and local nanomechanical indentation by applying *i-Rheo-Indent*, a recently introduced macroscale indentation methodology for the broadband rheological characterisation of soft solids [35], with an associated patent application [36]. Using a conventional tensile/compression testing machine equipped with a sensitive load cell, we apply the method to a 3D-printed liver analogue and to native murine quadriceps, brain, colon and liver tissues. The resulting viscoelastic spectra agree well with conventional oscillatory rheometry over the frequency range in which the latter remains reliable, while extending the experimentally accessible response up to 500 rad s^*−*1^, well beyond the high-frequency regime increasingly dominated by instrumental inertia in rotational measurements of these compliant samples. At the same time, *i-Rheo-Indent* retains the practical simplicity and geometrical flexibility of indentation, enabling measurements on intact and irregularly shaped specimens with minimal preparation. Thus, broadband tissue rheology can be performed using comparatively simple and accessible instrumentation while providing more reliable access to the high-frequency mechanical response.

## 2 Materials and Methods

### 2.1 Materials

In this work, validation is performed on several biologically relevant samples, which include a synthetic liver analogue material and murine tissues. The synthetic porcine liver analogue material, sourced from Stratasys Ltd., was 3-D printed using a Stratasys Digital Anatomy™ printer into an approximately cubic specimen (2 *×*2 *×*2 cm) according to the preset composition and architecture. Indentation measurements were first performed directly on these cubic specimens. Subsequently, the specimens were sectioned into cylindrical discs (15 mm in diameter and approximately 1.5 mm thick) for oscillatory rheometry.

Meanwhile, tissues were harvested from wild-type mice, with ages ranging between 8 to 16 weeks old. The collected tissue types comprised of brain (*n* = 3), quadriceps (*n* = 6), colon (*n* = 4), and liver specimens—three of which were kept intact and whole (*n* = 3) while another three were sectioned into 8-mm cylindrical disks (*n* = 3). Prior to testing, all murine tissues were briefly immersed in Hanks’ Balanced Salt solution (HBSS) prior to testing to prevent desiccation during transport and handling. Apart from sectioning the three liver specimens, no additional physical modifications were performed on the tissues prior to testing. All characterization was completed within a few hours of dissection, with oscillatory measurements performed prior to indentation testing.

### 2.2 Oscillatory Rheometry

Linear oscillatory measurements were performed on an Anton Paar MCR 302 rheometer using a 15-mm parallel-plate geometry for the synthetic sample and an 8-mm parallel-plate geometry for the murine tissue samples, as shown in Figure 1A. Amplitude sweep tests were first performed to identify the linear viscoelastic (LVE) regime of each material, as illustrated by the representative measurement in Figure 1B. Frequency sweep measurements were subsequently conducted within the identified LVE regime over the range 0.1–100 rad, s^*−*1^.

Within the framework of linear viscoelasticity, the measured shear stress *τ* (*t*) is related to the imposed shear history through the Boltzmann superposition principle [38]:

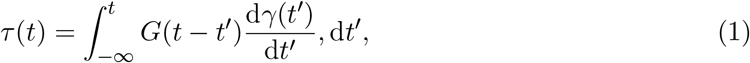

where *G*(*t*) is the shear relaxation modulus and *γ*(*t*) is the shear strain. Fourier transformation of this convolution relation yields the complex shear modulus [39]:

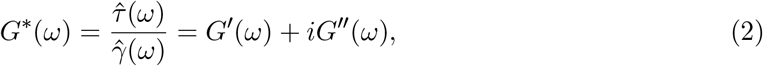

where *G*^*′*^(*ω*) and *G*^*′′*^(*ω*) are the storage and loss moduli, respectively, describing the elastic and dissipative components of the material response.

### 2.3 *i-Rheo-Indent* Measurements

All indentation measurements were conducted using a Mecmesin OmniTest 2.5 tensile/compression testing machine equipped with a 2-N load cell and a truncated-conical indenter featuring a 60^*°*^ cone angle and a 0.5-mm blunt-end diameter. Indentation depth was selected according to sample compliance, balancing the requirement to remain within the linear viscoelastic regime with the need to obtain a sufficiently resolved force response. An indentation depth of 100 *µ*m was used for most samples, unless specified otherwise, while 500 *µ*m was required for the biopsy-punched liver specimens. The resulting normal force was recorded for at least 100 s.

The experimental principle is illustrated in Figure 1D. A rigid truncated-conical indenter is brought into gentle contact with the sample and driven to a prescribed indentation depth, after which the displacement is held approximately constant while the resulting normal force relaxes in time. The indenter geometry is defined by its blunt-end radius *b* and cone angle *θ*, while the evolving contact radius is denoted by *a*. A representative measurement is shown in Figure 1E, where a rapid displacement to approximately 100 *µ*m produces a corresponding increase in normal force followed by progressive force relaxation.

The physical basis of the method follows directly from linear viscoelasticity and the Boltzmann superposition principle. Early work by Tripathy and Berger formulated stress-relaxation indentation by combining this principle with Hertzian contact mechanics, although the resulting relaxation response was subsequently interpreted by fitting prescribed viscoelastic models [40]. For an arbitrary indentation history *δ*(*t*), the measured normal force can be written in the general form:

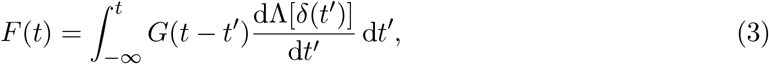

where Λ[*δ*(*t*)] is a geometry-dependent deformation function. F<u>o</u>r a rigid spherical indenter of radius *R*, classical Hertzian contact mechanics gives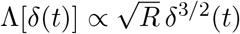, thereby incorporating the nonlinear relationship between indentation depth and contact area. In *i-Rheo-Indent*, a similar generalised formulation is applied to the truncated-conical geometry shown in Figure 1D, for which Λ[*δ*(*t*)] explicitly accounts for the indentation depth, blunt-end radius *b*, cone angle *θ*, and resulting contact radius *a* [35]:

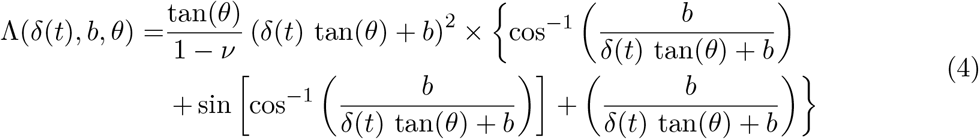

Thus, the contact mechanics are embedded directly into the deformation function, while the material response remains contained entirely in *G*(*t*). Fourier transformation converts Equation 3 into a simple product in the frequency domain, from which:

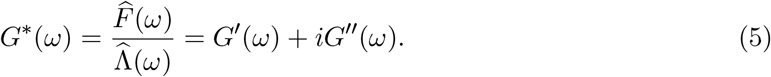

The experimentally measured force and indentation histories play roles analogous to stress and strain in conventional rheometry, while the contact geometry is explicitly embedded in Λ. Importantly, no ideal instantaneous step needs to be assumed: the actual time-dependent displacement imposed by the instrument is used in the transformation. Likewise, the viscoelastic response is recovered directly from the measured signals without fitting them to a predefined spring– dashpot constitutive model [39, 35]. Figure 1F shows the viscoelastic moduli obtained from the representative force-indentation curve in 1E.

The broadband nature of the measurement arises from the step-like excitation itself. In the ideal limit, the time derivative of a step is a Dirac impulse and therefore contains contributions over the full frequency spectrum. In practice, the experimentally accessible frequency window is determined by the shortest and longest resolved times of the measurement, while the measured displacement history accounts for the finite rise time of the imposed step. Consequently, a single force-relaxation experiment provides a continuous viscoelastic spectrum without requiring a sequence of oscillatory measurements at discrete frequencies.

This combination is particularly attractive for biological tissues. The macroscale contact area averages over a substantially larger volume than AFM, reducing sensitivity to microscopic heterogeneity, while indentation does not require the specimen to completely fill a predefined rheometer gap or to be machined into a regular disk. At the same time, because the broad-band response is obtained from relaxation rather than oscillatory motion, the high-frequency instrumental inertia that limits conventional shear rheometry is avoided. We therefore apply *i-Rheo-Indent* to synthetic tissue analogues and native murine tissues, including minimally manipulated specimens and samples with irregular geometries that are not readily accommodated by conventional parallel-plate rheometry.

## 3 Results and Discussion

The performance of *i-Rheo-Indent* was examined across three complementary experimental settings. First, measurements obtained from a synthetic liver analogue and murine quadriceps and brain tissues were compared with those from conventional oscillatory rheometry. The method was then applied to intact murine colon to demonstrate its suitability for irregular native geometries. Finally, intact and biopsy-punched murine livers were compared to assess how specimen preparation for conventional rheometry may influence the measured mechanical response.

### 3.1 Validation on Synthetic Porcine Liver Analogue Material and Murine Tissues

The first sample shown in Figure 2A is a 3-D printed porcine liver analogue material sourced from Stratasys Ltd., for which three replicates were measured using both *i-Rheo-Indent* and conventional oscillatory rheometry. During indentation measurements, a displacement of 100 *µ*m was applied. Overall, the two techniques showed good agreement across the frequency range in which the oscillatory measurements remained reliable. Because the synthetic material is considerably stiffer than the soft materials examined by Ewoldt *et al*. [20], the detrimental effects of instrument inertia were comparatively modest and became apparent only at relatively high frequencies, where a slight artificial upturn in the oscillatory response emerged. In contrast, *i-Rheo-Indent* continued to provide physically consistent spectra up to 500 rad, s^*−*1^. As shown below, the influence of instrumental inertia becomes substantially more pronounced for the softer native biological tissues.

**Figure 2:**
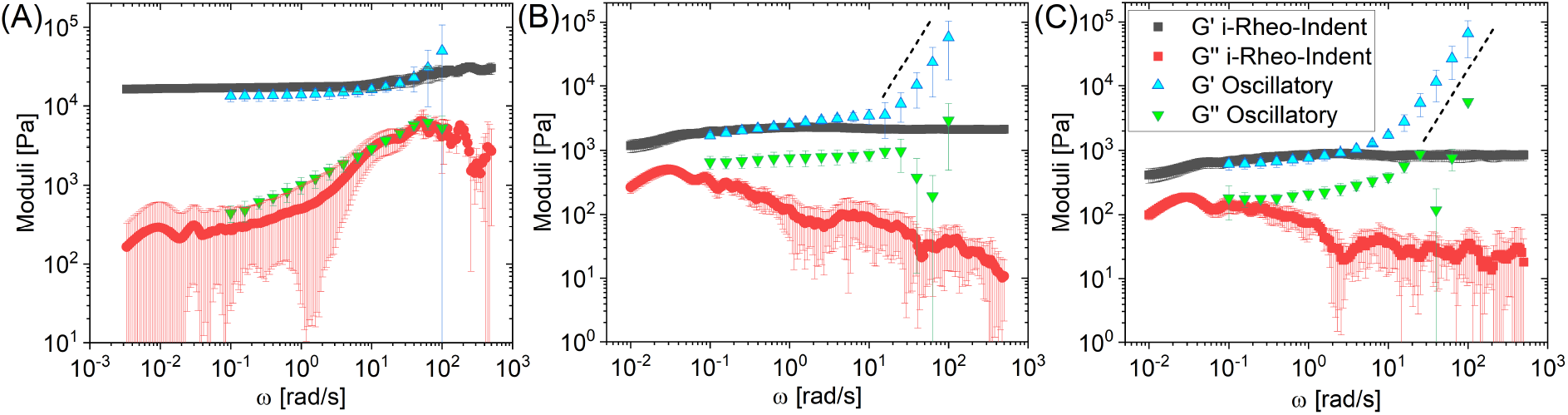
Comparison of viscoelastic moduli obtained via conventional oscillatory rheometry and *i-Rheo-Indent* for **(A)** a synthetic 3D-printed porcine liver phantom material, and wild-type murine tissues: **(B)** quadriceps and **(C)** brain. All three samples show good agreement with oscillatory measurements for the storage modulus *G*^*′*^, while *i-Rheo-Indent* circumvents the high-frequency inertial artifacts typically observed in oscillatory rheometry above *∼* 10 rad, s^*−*1^. The black dashed lines in (B) and (C) illustrate how instrument inertia creates apparent viscoelastic moduli in oscillatory measurements that scale almost to a factor of *ω*^2^.

Figure 2B presents the results obtained from quadriceps tissue, using two quadriceps harvested from each of three wild-type mice (*n* = 6), while Figure 2C shows the corresponding measurements for brain tissue (*n* = 3). Indentation measurements were performed three times per tissue sample, using indentation depths of 10 *µ*m for quadriceps and 100 *µ*m for brain, whereas oscillatory measurements were conducted once on each tissue sample. For each tissue type, indentation measurements were performed at comparable anatomical locations and with consistent specimen orientation across biological replicates; brain measurements were performed at comparable locations within the cerebrum. In contrast to the stiffer synthetic analogue, the substantially lower moduli of these native tissues make the detrimental effects of instrument inertia clearly apparent in the oscillatory measurements. At higher frequencies, the storage modulus *G*^*′*^(*ω*) exhibits a pronounced artificial upturn, approaching the characteristic *ω*^2^ scaling associated with instrument inertia, while the loss modulus *G*^*′′*^(*ω*) becomes increasingly unreliable and does not display a comparably clear trend. By contrast, *i-Rheo-Indent* continues to provide physically consistent spectra across the high-frequency regime, while remaining in good agreement with conventional oscillatory rheometry over the lower-frequency range in which the latter remains reliable.

For these samples, the loss modulus *G*^*′′*^(*ω*) exhibits visibly larger fluctuations than the storage modulus *G*^*′*^(*ω*). As discussed previously [35, 41, 42], these fluctuations arise from the propagation of experimental noise and finite acquisition resolution through the Fourier-transform procedure. This effect is particularly pronounced for predominantly elastic materials, for which *G*^*′*^(*ω*) *> G*^*′′*^(*ω*) and the smaller magnitude of *G*^*′′*^(*ω*) results in a lower signal-to-noise ratio. Consequently, the loss modulus is more susceptible to noise in the transformed spectra. Despite these fluctuations, the *i-Rheo-Indent* measurements remain in good agreement with oscillatory rheometry over the frequency range in which the latter remains reliable, particularly for *G*^*′*^(*ω*).

### 3.2 Murine Colons

We next applied *i-Rheo-Indent* to intact murine colon, a specimen with an irregular tubular geometry that is poorly suited to conventional parallel-plate rheometry. Figure 3 shows the viscoelastic moduli obtained from wild-type murine colon tissues (*n* = 4). Each colon was indented three times at comparable anatomical sites to a depth of 10 *µ*m, chosen to accommodate the thin and delicate nature of the tissue. This geometry highlights a key advantage of *i-Rheo-Indent*: mechanically relevant measurements can be performed directly on specimens in a near-native configuration, without reshaping them to fit a predefined rheometer geometry.

**Figure 3:**
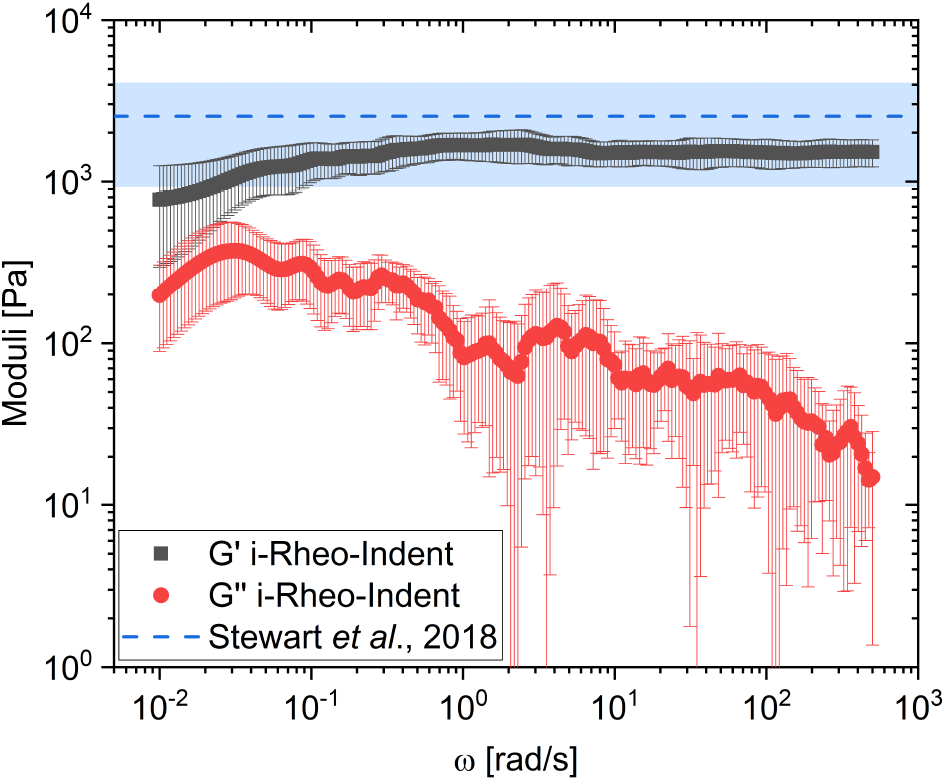
Comparison of the viscoelastic moduli obtained by *i-Rheo-Indent* for wild-type murine colon with the elastic modulus reported by Stewart *et al*. [43], rescaled to account for the different Poisson-ratio assumptions (*ν* = 0.4 in Stewart *et al*. and *ν* = 0.5 here) and subsequently converted to the equivalent shear modulus using *E* = 3*G*^*′*^.

In the absence of directly comparable oscillatory measurements, we compared our results with the murine colon stiffness reported by Stewart *et al*. [43], who performed indentation measurements using a spherical probe with a 1-mm diameter and analysed the response using a Winkler elastic-foundation model. Since their analysis assumed a Poisson’s ratio of *ν* = 0.4, whereas the present analysis assumes incompressibility (*ν* = 0.5), their reported elastic modulus (8.493 *±* 5.365 kPa) was first rescaled by the factor (1 *−*0.5^2^)*/*(1 *−*0.4^2^) to account for the different Poisson-ratio assumptions. The resulting value was then divided by 3, using *E* = 3*G*^*′*^ for an incompressible material, to obtain the equivalent shear modulus for comparison with our measurements. As shown in Figure 3, the elastic response measured by *i-Rheo-Indent* agrees well with the range obtained from the rescaled Stewart *et al*. data, despite the different indentation geometries and analytical approaches employed. Importantly, however, the Stewart *et al*. measurement provides only an elastic modulus, whereas *i-Rheo-Indent* additionally resolves the frequency-dependent viscous response through *G*^*′′*^(*ω*). Thus, while independently supporting the magnitude of the elastic response measured in intact colon, *i-Rheo-Indent* provides a more complete viscoelastic characterisation directly from the native, irregularly shaped tissue.

### 3.3 Intact and Cut Murine Livers

Finally, we examined whether the sample preparation required for conventional rheometry can influence the measured mechanical response of biological tissues. Livers were harvested from wild-type mice: three were measured whole and intact (*n* = 3), whereas three were cut into 8-mm cylindrical discs using a biopsy punch (*n* = 3) to match the parallel-plate geometry used for oscillatory rheometry. Indentation measurements were performed in triplicate for each sample, with representative force-relaxation and displacement histories shown in Figure 4A, while oscillatory measurements were conducted once per sample. The resulting viscoelastic spectra are presented in Figure 4B and C for the intact and cut livers, respectively.

**Figure 4:**
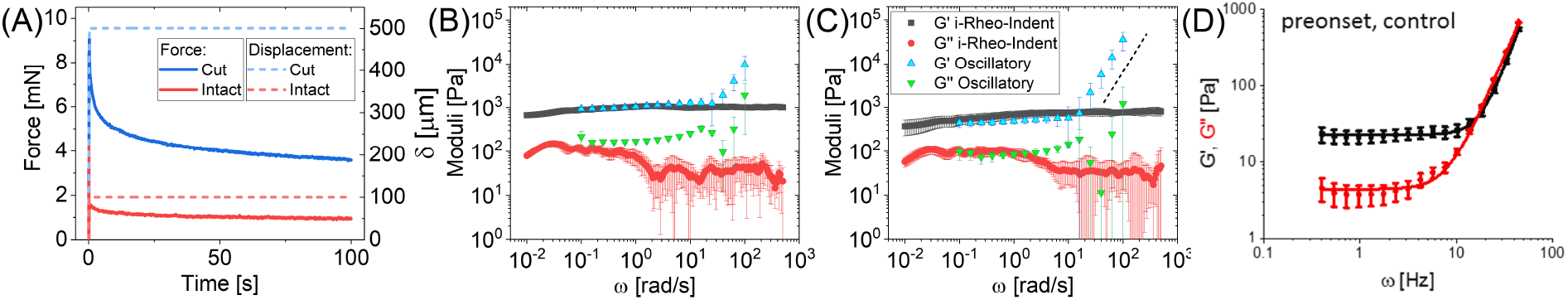
Comparison of viscoelastic moduli obtained from oscillatory shear rheometry and *i-Rheo-Indent* for intact livers and livers cut into 8-mm discs using a biopsy punch. **(A)** Representative force-relaxation and displacement histories for intact and biopsy-punched livers. An indentation depth of 100 *µ*m was applied to the intact livers, whereas 500 *µ*m was used for the biopsy-punched specimens to obtain agreement with the corresponding oscillatory rheometry measurements. **(B–C)** Frequency-dependent viscoelastic moduli recovered from the relaxation measurements in (A) for **(B)** intact and **(C)** biopsy-punched livers, plotted alongside conventional oscillatory rheometry measurements. The black dashed line in (C) indicates the approximate high-frequency scaling associated with instrumental-inertia artefacts in the oscillatory measurements. **(D)** Representative oscillatory frequency sweep for control murine liver reported by Pyka-Fościak *et al*. [44], showing the storage modulus *G*^*′*^ and loss modulus *G*^*′′*^ for the preonset control group. In that study, liver specimens were prepared as approximately 1-mm-thick, 4-mm-diameter discs after removal of the liver capsule, intrahepatic connective tissue and large blood vessels, and were subjected to 20% precompression prior to rheological testing. Reproduced from [44] with permission from Elsevier.

Most strikingly, the liver measurements reveal a pronounced dependence of the measured mechanical response on specimen preparation. The biopsy-punched liver samples exhibited substantially lower *G*^*′*^(*ω*) and *G*^*′′*^(*ω*) than the intact specimens, strongly suggesting that the manipulation required to accommodate biological tissue within a conventional rheometer geometry can materially influence the resulting mechanical characterisation. This observation is consistent with the substantially lower moduli reported by Pyka-Fościak *et al*. [44], who performed oscillatory rheometry on murine liver specimens across different stages of experimental autoimmune encephalomyelitis, with the baseline data from control (pre-onset) livers reproduced in Figure 4D. In that study, liver samples were frozen and subsequently thawed before being prepared as approximately 1-mm-thick, 4-mm-diameter discs, with the liver capsule, intrahepatic connective tissue, and large blood vessels excluded; a 20% precompression was then applied to ensure contact with the rheometer plate. Their reported storage modulus for control liver was 22.9 *±* 0.3 Pa, nearly two orders of magnitude lower than the values measured here for intact liver. Although the substantial differences in specimen preparation and experimental protocol preclude a direct quantitative comparison, the markedly lower modulus reported for these extensively prepared specimens further highlights tissue manipulation as a potential source of mechanical bias. Notably, the frequency sweeps reported by Pyka-Fościak *et al*. also display a pronounced high-frequency increase in both moduli, qualitatively consistent with the inertial artefacts observed in our conventional oscillatory measurements. This further illustrates the difficulty of accessing the high-frequency rheology of compliant tissues using rotational rheometry.

The effect of specimen preparation may be further amplified by the mechanics of the parallel-plate geometry itself. In parallel-plate shear, the local strain increases from zero at the centre to its maximum value at the outer radius [45]; consequently, the peripheral region—precisely where biopsy punching or trimming introduces the greatest structural disruption—is subjected to the largest deformation and contributes strongly to the measured torque. Consistent with this picture, an indentation depth of only 100 *µ*m was sufficient to obtain good agreement with oscillatory rheometry for the intact liver, whereas a substantially larger displacement of 500 *µ*m was required for the cut specimens. Such a fivefold increase in indentation depth is non-negligible for soft biological tissues, since larger deformations increase the risk of probing beyond the linear viscoelastic regime [46] and may make the measured response more sensitive to depth-dependent structural heterogeneity. Taken together, these observations indicate that specimen preparation should not be regarded as mechanically neutral: cutting or reshaping tissue to satisfy the requirements of conventional rheometer geometries may substantially influence the mechanical response ultimately attributed to the native tissue.

## 4 Conclusion

In conclusion, understanding how the mechanical properties of biological tissues regulate cellular behaviour, evolve with disease, and inform the design of biomaterials requires methods capable of resolving their frequency-dependent viscoelastic response while preserving, as far as possible, their native mechanical state. Here, we demonstrate that *i-Rheo-Indent* provides such a capability, recovering both *G*^*′*^(*ω*) and *G*^*′′*^(*ω*) from a single force-relaxation experiment while avoiding the high-frequency inertial limitations encountered in conventional oscillatory rheometry and substantially relaxing constraints on specimen geometry. Across synthetic and native soft tissues, the method showed good agreement with rotational rheometry over the frequency range in which the latter remained reliable, while extending measurements up to 500 rad, s^*−*1^. Importantly, it also enabled broadband viscoelastic characterisation of intact, irregularly shaped specimens such as murine colon, for which conventional parallel-plate rheometry would require substantial manipulation.

The liver measurements further highlight that specimen preparation is not necessarily mechanically neutral. Biopsy-punched specimens exhibited markedly lower viscoelastic moduli than intact livers, while literature data obtained from extensively prepared liver samples showed similarly low values, indicating that cutting, reshaping and compressing tissue to satisfy conventional rheometer geometries can substantially influence the mechanical response ultimately attributed to the native specimen. By combining rapid broadband acquisition, macroscale probing, minimal specimen manipulation and direct recovery of the complex viscoelastic response without fitting to a predefined constitutive model, *i-Rheo-Indent* provides a practical framework for characterising soft tissues closer to their native mechanical state. More broadly, this capability opens new opportunities for quantitative mechanobiology, mechanical phenotyping of healthy and diseased tissues, and the development and validation of tissue-engineered and biomimetic materials.

## Author Contributions

A.J.F. performed the indentation and rheological measurements, conducted the data analysis, and drafted the initial manuscript. L.J. harvested the murine tissue specimens and reviewed the manuscript. G.G. reviewed and edited the manuscript. M.T. conceptualized the study and revised the final manuscript.

## Acknowledgements

The authors would like to thank George Bruce for providing the synthetic liver analogue material and Collin Molloy for his assistance with the wild-type colons used in the study. A.J.F. additionally acknowledges financial support from the James Watt School of Engineering, University of Glasgow. M.T. acknowledges support from Innovate UK through the Knowledge Transfer Partnership programme (KTP-12905, 2021), supporting the collaboration between the University of Glasgow and Soapworks Ltd. The authors also gratefully acknowledge Soapworks Ltd for its continued support of the development of the *i-Rheo-Indent* methodology.

## Data Availability Statement

The data that support the findings of this study are available from the corresponding author upon reasonable request.

